# Interferon α and β induce differential transcriptional and functional metabolic phenotypes in human monocyte-derived macrophages and blunt glycolysis in response to antigenic stimuli

**DOI:** 10.1101/2023.12.16.571975

**Authors:** Gina Leisching, Anjali Yennemadi, Karl Gogan, Joseph Keane

**Affiliations:** TB Immunology Group, Department of Clinical Medicine, Trinity Translational Medicine Institute, School of Medicine, Trinity College Dublin, The University of Dublin, Dublin, Ireland

**Keywords:** type I interferons, metabolism, macrophage, glycolysis, oxidative phosphorylation, M. tuberculosis, LPS, transcriptomics

## Abstract

In the context of acute settings, the roles of type I interferons (IFNs), notably subtypes IFNα2a, 2b, and β, in modulating macrophage metabolism and contributing to host defense against viral and bacterial pathogens are well-established. However, the impact of chronic exposure to type I IFNs on macrophage metabolism, intimately linked to macrophage function, remains less understood. This study aimed to unravel the nuanced host responses induced by type I IFN cytokines, offering insights for potential therapeutic approaches in diseases associated with these cytokines.Employing a combination of transcriptional profiling and real-time functional analysis, we delineated the temporal evolution of metabolic reprogramming in response to chronic interferon exposure. Our results reveal distinct transcriptional metabolic profiles between macrophages chronically exposed to IFNα and IFNβ. Agilent Seahorse assays demonstrated that IFNβ significantly diminishes the oxygen consumption rate and glycolytic proton extrusion rate in macrophages. Conversely, IFNα2b decreased parameters of mitochondrial fitness and induced a shift towards glutamine oxidation.Assessing the ability of macrophages to induce glycolysis in response to antigenic stimuli (LPS and iH37Rv), we found that chronic exposure to all IFN subtypes limited glycolytic induction. This study addresses a critical oversight in the literature, where individual roles of IFN subtypes are frequently amalgamated and lack distinction. These findings not only provide novel insights into the divergent effects of interferon α2a, α2b, and β on macrophage metabolism but also highlight their potential implications for developing targeted therapeutic strategies. This is particularly relevant in autoimmune disorders where type I IFNs, particularly IFNα, play a central role. The observed metabolic quiescence induced by chronic IFN exposure underscores its significance in macrophage functionality and its potential contribution to the pathophysiology of autoimmune disorders and susceptibility to infection.

## Introduction

Cellular metabolism plays a critical role in regulating and fine-tuning immune function where it serves as a pivotal orchestrator of immune responses. Macrophages, key players in the innate immune system, undergo dynamic metabolic alterations upon stimulation with type I interferons (IFNs).^1^ Acute exposure to type I IFNs has been previously demonstrated to elicit metabolic shifts in macrophages, including increased glycolytic flux, inhibition of sterol biosynthesis, redirection of lipid metabolism from de novo synthesis to lipid import, and enhanced tryptophan catabolism.^2^

In the context of autoimmune diseases, such as systemic lupus erythematosus (SLE), where chronic type I IFN secretion is a hallmark, macrophages exhibit profound hypermetabolic phenotypes that correlate with disease severity.^3,4^ Despite the wealth of knowledge regarding acute responses, there remains a critical gap in understanding the metabolic reprogramming associated with chronic type I IFN exposure, particularly concerning the nuanced differences between distinct interferon subtypes.

To address this gap, the present study focuses on systematically characterizing the metabolic consequences of chronic type I IFN responses in human monocyte-derived macrophages (hMDMs). Specifically, we investigate the impact of chronic stimulation with three representative type I IFNs—α2a, α2b, and β—on the global metabolic gene expression profile of hMDMs. Notably, these interferon subtypes are thought to exert unique functional effects in various cellular contexts. Further, we aimed to test the metabolic functional response of the treated macrophages to induce glycolysis following stimulation with the antigens LPS and irradiated H37Rv, since this is crucial in the host defence against infection.^5,6^

The rationale behind this study is rooted in the need to unravel the intricate metabolic signatures induced by different type I IFN subtypes, despite their shared classification. By employing transcriptional profiling and real-time functional analysis, we aim to delineate the temporal evolution of metabolic reprogramming in response to chronic interferon exposure. Our findings are poised to provide novel insights into the divergent effects of interferon α2a, α2b, and β on macrophage metabolism, paving the way for a clearer understanding of the immunometabolic landscape in the context of chronic macrophage activation. Ultimately, such insights hold potential implications for developing targeted therapeutic strategies in autoimmune disorders where type I IFNs play a central role.

## Materials and Methods

### Cell culture

Peripheral blood mononuclear cells (PBMC) were isolated from the buffy coats of healthy donors, obtained with consent from the Irish Blood Transfusion Services, by density-gradient centrifugation over Lymphoprep (StemCell Technologies). Cells were washed, resuspended at 2.5x10^6^ PBMC/ml in RPMI (Gibco) supplemented with 10% AB human serum (Sigma-Aldrich), and plated onto non-treated tissue culture plates (Costar). Cells were maintained in humidified incubators for 7 days at 37°C and 5% CO2. Non-adherent cells were removed by washing every 2-3 days. The purities of MDM were assessed by flow cytometry and were routinely > 90% pure.

### Chronic type I IFN exposure (CT1IE)

Human IFNβ (IF014, Merck, KGaA, Darmstadt), IFNα2a (11100-1, PBL assay science, NJ, USA), and IFN IFNα2b (11105-1, PBL assay science, NJ, USA) concentrations were determined through a dose response curve, where expression of ISG15 was used as a marker for activation of the type I IFN pathway (Supplementary figure 1). As described previously, IFN α2a, α2b, and β were added at a concentration of 1000U/ml for 5 days before analysis.^7^ This concentration of IFNs increased ISG expression to the same degree seen in IFN-High SLE patients and was therefore used in all subsequent experiments. No evidence of cell death was detected after the 5-day period (Supplementary Figure 1).

### RNA extraction and Nanostring Technology

On day 5 of CT1E, RNA from each treatment group was extracted using the RNeasy® Plus Mini Kit (Cat. No. 74134, Qiagen, Limburg, Netherlands) according to the manufacturer’s instructions. The ‘gDNA eliminator’ column included in this kit removed genomic DNA in all samples. For each experiment, RNA quality and quantity was assessed and only RNA with a RNA integrity Number (RIN) above 9.0 was used for Nanostring analysis. The NanoString nCounter Human Metabolic Pathways Panel (Cat # XT-CSO-HMP1-12, N = 6 per IFN sub-type), which detects transcripts of 768 metabolic genes. All data were analyzed using nSolver 4.0. Background was subtracted from negative controls and samples were normalized to the positive controls and housekeeping genes. Data were further analyzed by normalized count and fold difference, and differences were compared between the sub-types.

### Metabolic Assays Using the Seahorse XFe Analyzer

Macrophage metabolic function was assessed using the Seahorse XFe Analyzer (Agilent) where mitochondrial function (Mitostress Test), mitochondrial fuel usage (MitoFlex fuel test), and glycolytic rate (Glycolytic rate assay) were assessed post-IFN treatment using a Seahorse XF extracellular flux analyser (Seahorse Bioscience, Inc, North Billerica, MA, USA). For the induced glycolytic assay, either LPS (10ng/ml) or irradiated H37Rv (iH37Rv) with a predetermined multiplicity of infection (MOI) was added into ‘Port A’ of the Agilent cartridge. Rotenone and 2-DG were then added sequentially into port B and C. The same GlycoPER values for ‘untreated’, ‘LPS’ and ‘iH37Rv’ are used within the separate graphs representing IFNα2a, 2b, and β in Figure 4D as all experiments were run using matched donors. All values were normalised using the Crystal Violet dye extraction growth assay and the Wave Desktop 2.6 Software (https://www.agilent.com) was used for analysis.

### qPCR

Total RNA was extracted using the RNeasy® Plus Mini Kit (Cat. No. 74134, Qiagen, Limburg, Netherlands) according to the manufacturer’s instructions immediately after the treatment period. The “gDNA eliminator” column removed genomic DNA in all samples. For cDNA synthesis, 0.5 µg RNA was converted to cDNA using the Quantitect® Reverse Transcription Kit (Cat. No. 205311, Qiagen, Limburg, Netherlands). gDNA wipe-out buffer was added to RNA prior to the RNA conversion step. qPCR amplification was performed in 96-well plates and run on a QuantStudio5 (Thermofisher Scientific). LightCycler® 480 SYBR Green I Master (Cat. No. 04887352001, Roche, Germany) was used for various differentially expressed genes (*HK2, PFKFB3, PKM2*) using QuantiTect® primer assays at a reaction volume of 20 µl with reference genes *UBC* and *16S*. The amplification procedure entailed 45 cycles of 95°C for 10 s followed by 60°C for 10s and finally 72°C for 10s. Fold change in expression levels calibrated using the 2 -ΔΔCT method. All biological replicates were run in triplicate with a positive control (calibrator) and a non-reverse transcription control in accordance with the MIQE Guidelines.

### Evaluation of mitochondrial membrane potential via JC-1 immunofluorescent staining

JC-1 can detect mitochondrial membrane potential as a probe. It aggregates in the matrix at high membrane potential and emits red fluorescence (590 nm). However, the JC-1 keeps acting as a monomer at low membrane potential and emits green fluorescence (525 nm).Human monocyte-derived macrophages (hMDM) were seeded onto glass coverslips at a concentration of 2x 10^6^ cells/ml in a 24 well plate and allowed 7-10 days to differentiate. After CT1IE, macrophages were treated with JC-1 working solution (1 μg/ml) for 25 minutes at 37 °C. Cells were imaged and captured at ×63 magnification using a Leica SP8 scanning confocal microscope. Confocal images were quantified by calculating the ratio of mean red fluorescence/green fluorescence of each cell using Image J software. Four fields of view per image, with 3 images per condition within 3 replicates (donors) were assessed.

### Statistical analysis

Statistical analyses were performed using GraphPad Prism 9 software. Statistically significant differences between three or more groups were determined by one-way ANOVA with Tukey’s multiple comparisons tests. P-values of <0.05 were considered statistically significant.

## Results

### IFNs α and β induce transcriptionally distinct metabolic profiles in human MDMs

Firstly, we employed transcriptomics to elucidate the intricate interplay between the interferon subtypes and the host macrophage metabolism over time. This approach allowed us to uncover potential subtype-specific effects and temporal dynamics in the regulation of metabolic genes. As shown in the PCA plot (Fig. 1A), the untreated macrophages (Control) form a discrete cluster, indicating a distinct transcriptional profile associated with the baseline state of macrophages in the absence of interferon stimulation. Notably, macrophages treated with IFN-β exhibit a separate and clearly defined cluster, suggesting a unique transcriptional response to interferon beta stimulation. Unsurprisingly, the PCA plot reveals that macrophages treated with IFN-α2a and IFN-α2b cluster together, indicating a similar transcriptional signature induced by these two interferon subtypes. The clustering of IFNα2a and IFNα2b is consistent with their shared classification within the type I interferon family. The distinct separation of IFNβ-treated macrophages from the IFN-α2a and IFN-α2b cluster aligns with the notion that different interferon subtypes can elicit unique transcriptional responses in macrophages. This finding underscores the importance of exploring subtype-specific effects to unravel differences in macrophage interferon signaling. The Venn diagram (Fig1.B) shows that 21.9% of the metabolic genes induced are shared by all three subtypes. Macrophages treated with IFNα2a express 3.8% of unique genes, whereas IFNα2b-treated macrophages only express 1.9% of unique genes. Interestingly, macrophages treated with IFNβ express 62.1% of genes that are not shared with any other subtypes. Volcano plots depict up- and downregulated metabolic genes in macrophages treated with each subtype compared to the control. *STAT1, GMPR*, and *TYMP* were the most up-regulated genes across all subtypes, and *RPLP0* was the most downregulated gene across all subtypes.

**Figure 1.**
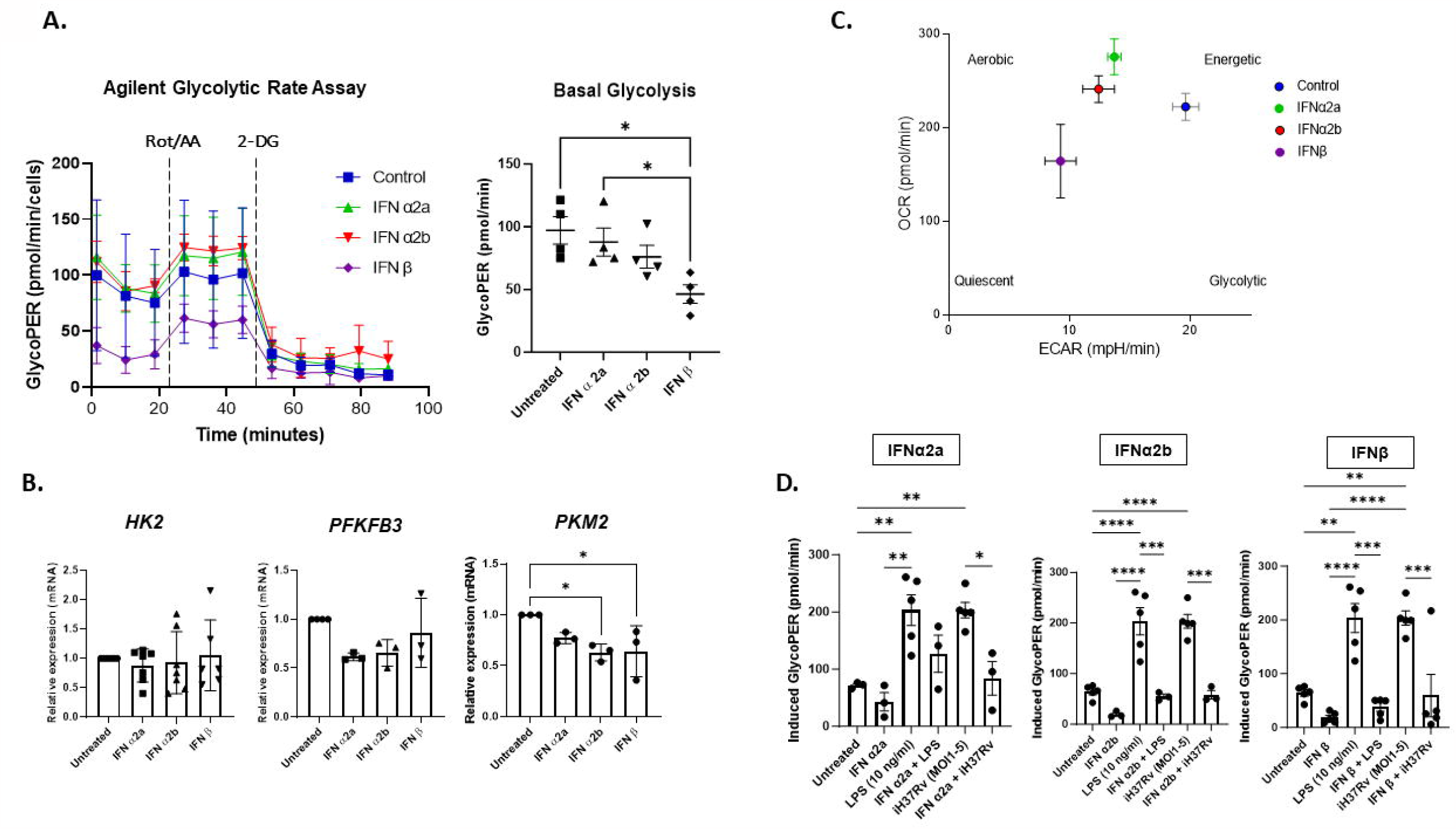
IFNs α2a, α2b, and β induce transcriptionally distinct metabolic profiles in macrophages. **A**. Principal Component Analysis (PCA) plot illustrating the distinct clustering patterns of untreated macrophages (Control), macrophages treated with interferon beta (IFN-β), and those treated with interferon alpha 2a and 2b (IFNα2a and IFNα2b). **B**. Venn diagram indicating shared and unique metabolic genes between the IFN subtypes. **C**. Heatmap displaying each sample’s directed global significance scores for each metabolic pathway. Directed global significance statistics measure the extent to which a gene set’s genes are up- or down-regulated with the variable. Red denotes gene sets whose genes exhibit extensive over-expression with the covariate, blue denotes gene sets with extensive under-expression. **D**. Volcano plots indicating both up- and down-regulated genes in macrophages in response to each IFN subtype, depicted as log2(fold change) vs. untreated control.

### IFNβ significantly reduces basal oxygen consumption rate (OCR) and mitochondrial fitness in human MDMs

To determine whether CT1IE affected mitochondrial respiratory capacity, we used the Agilent Mitostress test (Fig.2). We observed typical Mitostress profiles in macrophages treated with all IFN subtypes (Fig.2A), however, IFNβ significantly reduced basal OCR levels in comparison to untreated macrophages. Maximal respiration has significantly reduced the treatment of macrophages with IFNα2b and IFNβ vs. control. Interestingly, macrophages treated with IFNα2a had no significant effect on maximal respiration. Next, we assessed the spare respiratory capacity and observed that treatment with IFNα2a had significantly higher spare respiratory capacity compared to macrophages treated with both IFNα2b and IFNβ. Next, we wanted to determine whether mitochondrial membrane potential(MMP) was affected in macrophages treated with the IFN subtypes(Fig. 2B). Surprisingly, macrophages treated with IFNα2a had a higher MMP than macrophages treated with both IFNα2b and IFNβ.

**Figure 2.**
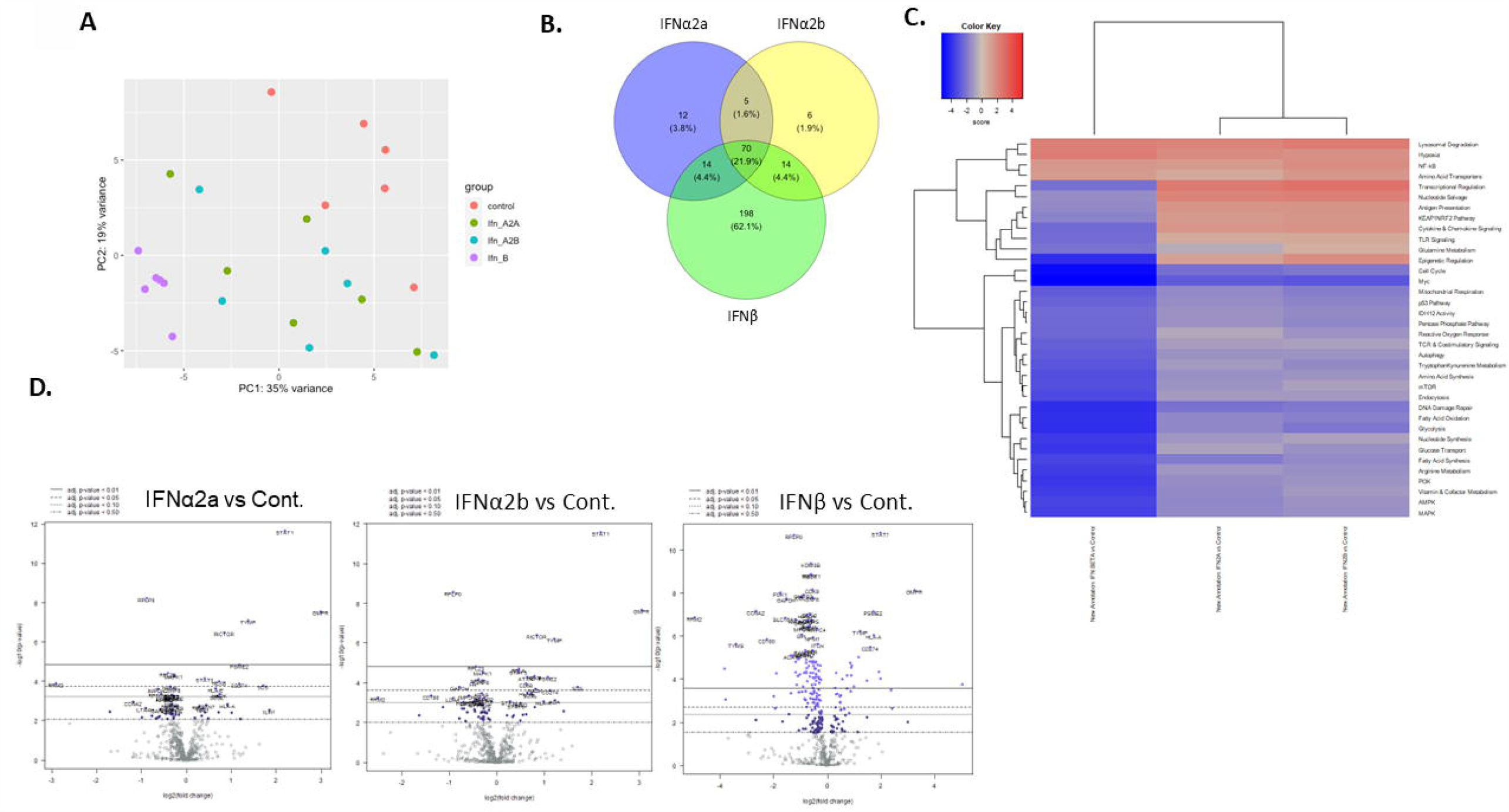
Macrophages chronically exposed to IFNβ have significantly reduced basal OCR and mitochondrial fitness. A. The Agilent Mitostress test was used to assess mitochondrial fitness and function during sequential treatment with oligomycin, carbonyl cyanide-4-(trifluoromethoxy)phenylhydrazone (FCCP) and antimycin A + rotenone. IFNβ induced significant reductions in basal OCR, maximal respiration, and spare respiratory capacity vs. untreated control and IFNα2a treated macrophages. Maximal respiration was also significantly impaired in macrophages treated with IFNα2b vs. untreated control. Spare respiratory capacity was significantly increased with macrophages treated with IFNα2a vs. IFNα2b and IFNβ. Data shown are the mean ± SEM (n = 4-7). One-way ANOVA with Multiple comparisons test ^*^ p < 0.05, ^**^ p < 0.01; ^***^ p < 0.001. Scale bar = 10 μm.

### Macrophages undergo mitochondrial metabolic modifications after treatment with IFNα2a, IFNα2b and IFNβ

Since we observed varying changes in mitochondrial function, we next sought to determine whether substrate oxidation and preference were affected after CT1IE with the IFN subtypes. We employed the Mito Fuel Flex Test to measure fuel pathway preference by sequentially modulating glucose, glutamine and fatty acid oxidation using the inhibitors UK5099, BPTES and Etomoxir respectively (Fig.3A). Untreated macrophages show dependence on all three fuels at similar levels of dependency (Fig.3B). Macrophages treated with IFNα2a show an increased dependency on glucose oxidation, whereas macrophages treated with IFNα2b have a higher dependency on glutamine oxidation in comparison to control. Conversely, macrophages treated with IFNβ show reduced dependency on glucose and glutamine oxidation when compared with macrophages treated with the other two subtypes, however, these effects were not significant when compared to untreated macrophages. Macrophages treated with all IFN subtypes did not show changes in the dependency for fatty acids as a fuel source. To probe these findings further, qPCR to detect transcription levels of the three fuel substrate transporter/enzyme genes (*MPC, CPT1*, and *GLS*) was assessed (Fig. 3C). mRNA levels of mitochondrial pyruvate carrier (*MPC*) were significantly decreased in macrophages treated with both IFNα2b and IFNβ in comparison to macrophages treated with IFNα2a. *MPC* expression was upregulated in macrophages treated with IFNα2a and agrees with Mitostress results showing an increase in glucose oxidation. Glutaminase (*GLS*) mRNA was significantly increased in macrophages treated with IFNα2b compared to control and is in line with functional data (Fig.3B) but remain unchanged in macrophages treated with IFNα2b and IFNβ. Carnitine palmitoyltransferase 1 (*CPT1*) mRNA was significantly downregulated in macrophages treated with all three subtypes. Despite this finding, oxidation of fatty acids remained unchanged (Fig. 3B).

**Figure 3.**
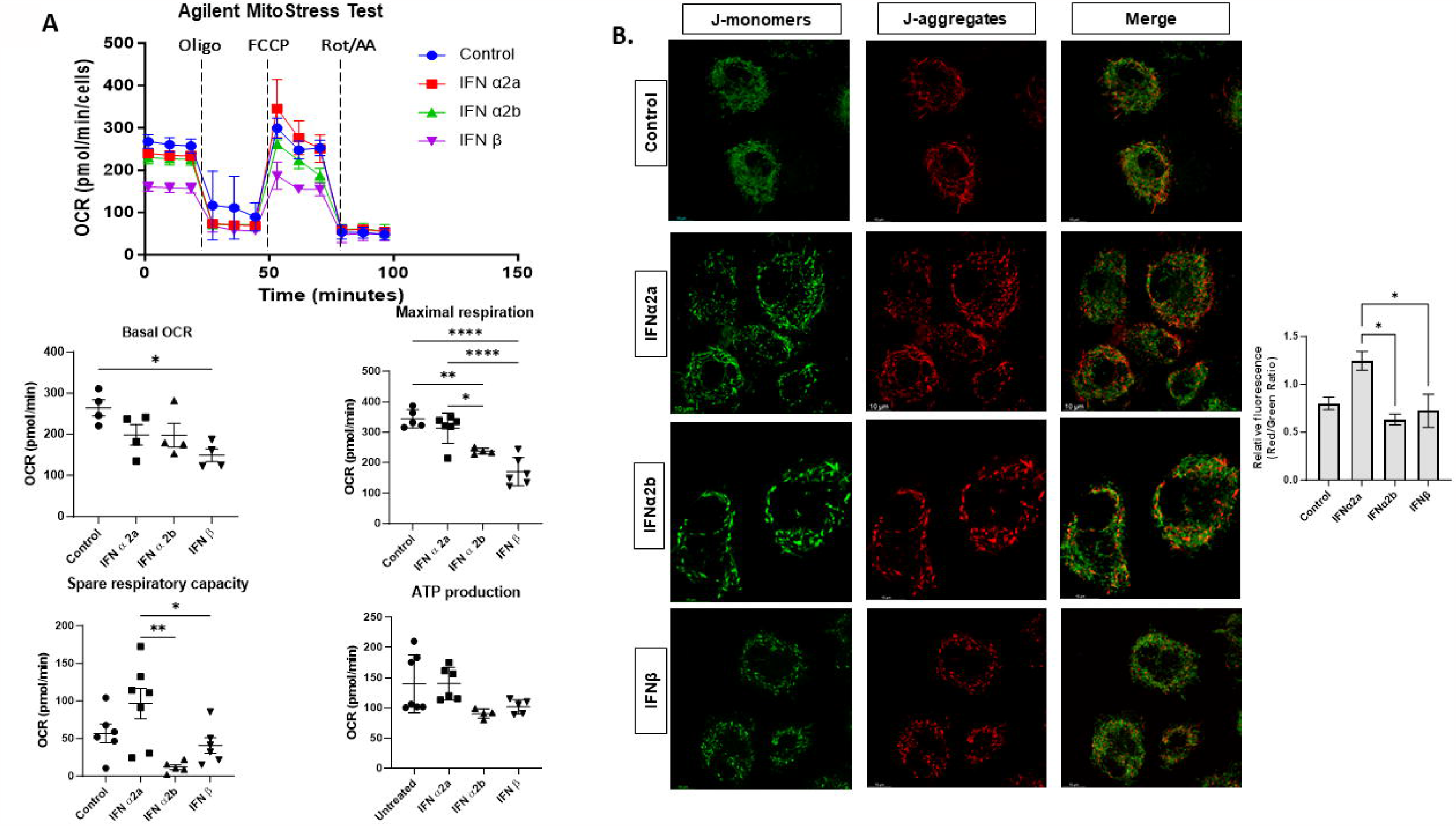
IFNα2a and IFNα2b treatment induces a shift toward glucose and glutamine dependence in macrophages. **A**. The Mito fuel flex test inhibits the importation of three major metabolic substrates (pyruvate, fatty acids, and/or glutamine) with mitochondrial pyruvate carrier inhibitor UK5099 (2 mM), carnitine palmitoyltransferase 1A inhibitor etomoxir (4 mM), or glutaminase inhibitor BPTES (3 mM), figure adapted and modified from https://www.agilent.com/cs/library/usermanuals/public/XFp_Mito_Fuel_Flex_Test_Kit_User_Guide.pdf **B**. Mitochondrial fuel dependency tests show that FAO remains unchanged, however, macrophages treated with IFNα2a significantly increase glucose oxidation, whereas macrophages treated with IFNα2b have an increased dependence on glutamine oxidation. Treatment with IFNβ decreased oxidation of both glucose and glutamine vs. IFNα2a and 2b. **C**. qPCR too assess gene expression of *MPC, GLS*, and *CPT1* to confirm findings from Mito flex fuel data. Macrophages treated with IFNα2a significantly increase MPC gene expression, whereas macrophages treated with IFNα2b significantly increase *GLS* expression. Treatment with all subtypes decreases *CPT1* expression. Data shown are the mean ± SEM (n = 3-7). One or two-way ANOVA with Multiple comparisons test ^*^ p < 0.05, ^**^ p < 0.01; ^***^ p < 0.001.

### Glycolysis is significantly reduced in macrophages following treatment with IFNβ

Next, we wanted to determine whether glycolysis was affected by CT1IE (Fig. 4). Glycolytic Rate Assay utilizes both extracellular acidification rate (ECAR) and OCR measurements to determine the glycolytic proton efflux rate (glycoPER) of the cells. GlycoPER is the rate of protons extruded into the extracellular medium during glycolysis as well as compensatory glycolysis following mitochondrial inhibition. Macrophages treated with IFNβ have significantly lower glycoPER in comparison to control and IFNα2a treated macrophages (Fig. 4.A). No differences in glycoPER were observed in macrophages treated with IFNα2a or 2b when compared to the control. We were then interested in assessing the expression of three glycolytic genes, hexokinase (*HK2*), 6-phosphofructo-2-kinase (*PFKFB3*), and pyruvate kinase (*PKM2*) which represent the beginning, middle, and end of the glycolytic pathway, respectively. Our results show that only *PKM2* expression was significantly decreased in macrophages treated with IFNα2b and IFNβ in comparison to control (Fig. 4.B), suggesting that the reduction in glycolysis may in part be due to decreased PKM2 expression. Finally, we combined both OCR and GlycoPER readings to generate an energy metabolism phenotype representing macrophages treated with all IFN subtypes (Fig.4C). Unsurprisingly, macrophages treated with IFNα2a and 2b clustered together, depicting a metabolic phenotype that is more aerobic in comparison to control, whereas macrophages treated with IFNβ exhibited a less metabolic phenotype, possibly moving toward metabolic quiescence.

**Figure 4.**
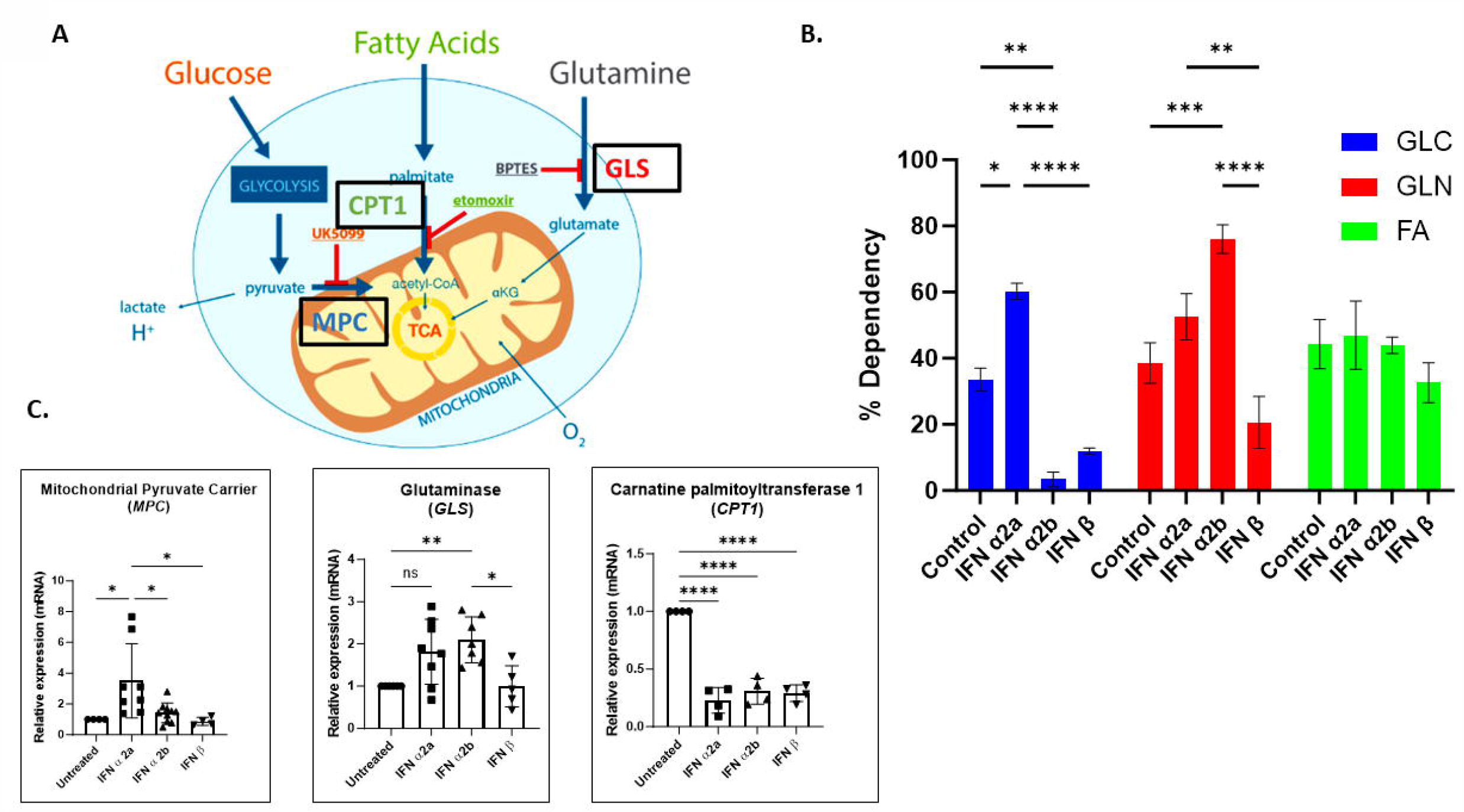
Macrophages chronically exposed to IFNβ have significantly reduced basal GlycoPER and *PKM2* expression and fail to induce glycolysis in response to acute LPS or iH37Rv stimulation. **A**. GlycoPER was assessed using the Agilent glycolytic rate assay where ECAR baseline readings were recorded using the Seahorse XFp analyzer, whereafter 4 µM Rot/AA and 50 mM 2-deoxyglucose (2-DG) were injected respectively. Basal glycolysis was significantly reduced in macrophages chronically treated with IFNβ vs. untreated and IFNα2a-treated macrophages. **B**. qPCR showing gene expression of three glycolytic enzymes, *HK2, PFKFB3*, and *PKM2* following treatment. PKM2 expression was significantly reduced in macrophages chronically treated with IFNα2b and IFNβ vs. untreated control. **C**. Metabolic phenogram of baseline readings depicting macrophages treated with IFNα2a and 2b clustering together, and macrophages treated with IFNβ depicting a less aerobic and energetic phenotype. **D**. The induced glycolytic assay was conducted to determine the ability of macrophages to respond to antigenic stimulation with LPS and iH37Rv. Untreated macrophages induce a significant induction of glycolysis 20 min after the injection of the stimuli into the wells, however, macrophages treated with all subtypes were unable to significantly induce glycolysis compared to these controls. Data shown are the mean ± SEM (n = 3-5). One or two-way ANOVA with Multiple comparisons test ^*^ p < 0.05, ^**^ p < 0.01; ^***^ p < 0.001.

Finally, we wanted to test the ability of these macrophages to respond appropriately to antigenic stimuli using two well-known macrophage-stimulating bacterial components, LPS and iH37Rv. Normally, macrophages rapidly upregulate glycolysis in response to these stimuli which is linked to the production of pro-inflammatory mediators to resolve the infection.^5,8,9^ To test the induction of glycolysis, we used the induced Glycolytic Rate Assay that allows for real-time quantification of *in situ* glycolysis activation or repression after a stimulus (Figure 4D). As expected, stimulation of untreated macrophages with LPS or iH37Rv alone significantly induced glycolysis, however, macrophages chronically exposed to either IFNα2a, IFNα2b or IFNβ were unable to induce glycolysis when stimulated with these antigens. Thus, type I IFNs significantly hinder the macrophages glycolytic capacity.

## Discussion

Type I interferons (IFN) can have dual and opposing roles in immunity, with effects that are beneficial or detrimental to the individual depending on whether IFN pathway activation is transient or sustained. This study addresses a prevailing oversight in the literature where the individual roles of IFN subtypes are frequently amalgamated, lacking distinction. Our findings demonstrate that human monocyte-derived macrophages exhibit unique and subtype-specific metabolic responses to IFN-α2a, IFN-α2b, and IFN-β. Notably, chronic exposure to these IFN subtypes is shown to impede macrophage glycolysis in response to antigenic stimuli, highlighting the need for a nuanced understanding of the immunometabolic consequences of distinct type I IFN subtypes in immune regulation.

In the present study, prolonged exogenous administration of different interferon (IFN) subtypes to human monocyte-derived macrophages revealed distinct metabolic transcriptomes induced by IFNα2a, IFNα2b, and IFNβ. Notably, our findings demonstrate a significant downregulation of genes involved in glycolysis and mitochondrial respiration, presenting a stark contrast to transcriptional studies focused on acute macrophage stimulation, particularly with IFNα,^10^ which typically show an overall increased metabolic shift. This dichotomy underscores the complexity of the temporal dynamics in macrophage responses to IFNs and emphasizes the need for careful interpretation of IFN subtype-specific effects. Importantly, our observation of a sustained decrease in glycolytic and mitochondrial gene expression highlights the biological significance of chronic type I IFN exposure. This metabolic quiescence induced by prolonged IFN stimulation bears particular relevance in diseases characterized by persistent type I IFN responses, such as SLE, where macrophage functionality may be compromised thus contributing to the overall pathophysiology.^11^

Furthermore, our functional analysis shows that IFNβ significantly reduces mitochondrial fitness and basal glycolysis. Surprisingly, mitochondrial fitness was least affected by chronic exposure to IFNα2a. When assessing mitochondrial fuel usage, our work showed no difference in FA oxidation (FAO). This finding is in contrast to a study by Wu and colleagues that showed increased FAO and oxidative phosphorylation (OXPHOS) following IFNα treatment.^12^ The difference in these findings may be attributed to the fact that peripheral dendritic cells(pDCs) were used in the reported study and only 24 h of exposure to IFNα was conducted. Thus, the observed alterations in FAO in pDCs may therefore be an acute response to IFNα exposure. Results in line with our findings are presented in a study that includes a chronic IFNα exposure model on CD8 T cells.^7^ The authors found that CD8 T cells from SLE patients with high IFN-stimulated genes (ISGs) signatures had a downregulation of OXPHOS genes as well as significantly lower basal and maximal OCR and spare respiratory capacity, increasing cell death. Thus, prolonged exposure to both IFNα and IFNβ induces metabolic dysregulation that may contribute to pathogenesis.

This study has shown clear differences between the metabolic profile of macrophages treated with IFNα and β, however, the differences between the two IFNα subtypes are more subtle. This was not surprising since therapy with both peginterferon-α-2a and peginterferon-α-2b appear from comparative studies to be similarly tolerated, with few differences of clinical significance noted.^13^ The most noteworthy difference is their mitochondrial fuel preference. The preferential induction or shift towards glutamine oxidation by macrophages chronically exposed to IFNα2b, and glucose oxidation by macrophages exposed to IFNα2a was unexpected. Several publications report increased glutamine/glutamate levels after IFNα therapy^14,15^ however the authors fail to report the IFN subtype used in these studies. Glutamine utilization has been linked to functional activities of macrophage function such as cytokine production, nitric oxide production, superoxide production, and phagocytosis.^16^ Assessing these parameters will provide valuable insights into the broader implications and functional consequences of the various IFN subtypes on macrophage functionality and its role in disease.

The type I IFN cytokine family encompasses over 10 IFNα members and a single IFNβ, signals through the IFNAR receptor, inducing the modulation of hundreds of genes. While this response is pivotal for establishing an antiviral state, the regulatory role of type I IFN in immune responses to intracellular bacterial pathogens remains a subject of ongoing investigation. The impact of IFNα/β on host outcomes during bacterial infections is dichotomous, with both protective and detrimental effects observed. The precise mechanisms underlying the exacerbation of disease driven by IFNα/β are complex and not fully elucidated. Initial investigations into hyper-virulent M. tuberculosis strains suggested that the suppression of pro-inflammatory cytokines and TH1-type immunity is significant. ^17^ Notably, evidence from studies in human cells and mouse models indicates that IFNα/β may suppress the production of host-protective cytokines following *M. tuberculosis* infection, contributing to the multifactorial nature of this regulatory phenomenon. The production of IL-1β, a pro-inflammatory cytokine that is dependent on glycolysis and is crucial for host defense against M. tuberculosis, is inhibited by IFNα/β, both in vitro in infected human and mouse cells and in vivo.^18,19^ A recent study found that type I IFNs, specifically IFNβ, decrease macrophage energy metabolism during mycobacterial infection.^20^ Employing our in vivo model of chronic IFNα2a treatment in male Wistar rats, we stimulated bone marrow-derived macrophages (BMDMs) with iH37Rv, ^21^ replicating the experimental approach described in this paper. Notably, we observed that BMDMs from rats treated with IFNα2a were less effective in upregulating glycolysis in response to iH37Rv compared to BMDMs from rats treated with saline. This consistent observation aligns with the results presented in this study and those described earlier, reinforcing the impact of both IFNα and IFNβ exposure on the glycolytic reprogramming of macrophages to antigenic stimuli, and therefore the macrophage host response mechanism. The results presented in this study only account for approximately 20 min post-stimulation. The effects of the various subtypes on longer infection times and therefore infection outcome is an important future study that should be conducted.

In conclusion, this study illuminates the intricate interplay between type I IFNs and macrophage metabolism, offering a foundation for future investigations that aim to unravel the nuanced roles of individual IFN subtypes and their broader implications in immune regulation and disease pathogenesis.

## Supplementary Figure Legends

**Figure S1.IFNα and β dose-response and cell viability assessment in human macrophages chronically treated with either IFNα2a, 2b or β for 5 days.** A. Representative Western blots to determine ISG15 expression as a marker of IFN pathway induction. 1000IU/ml was chosen for all experiments. B. Cell viability assays using PI as a marker of cell death with corresponding images taken on the light microscope. No significant differences in cell death were observed following treatment. Scale bar = 100 uM.

