## Supplemental methods for "Interferon α and β induce differential transcriptional and functional metabolic phenotypes in human monocyte-derived macrophages and blunt glycolysis in response to antigenic stimuli"

**Supplementary Methods**

Western Blotting

Macrophages were lysed with RIPA buffer and whole protein extracts quantified using the CB-X proein assay(G-Biosciences) and were then analysed through Western blotting. Membranes were incubated with the primary antibodies (1:1000 dilution) β-actin (loading control) and ISG15 (Cell Signaling, MA, USA). Membranes were then washed and incubated with anti-rabbit secondary antibodies (1:5000) for 1h and then exposed using the ChemiDoc MP Imaging System (BioRad) after the addition of ECL for 1 min.

Cell viability assessment

After the treatment period, the macrophages were stained with PI (2 mg/ml) and DAPI and then examined and quantified with a Lionheart FX automated microscope.
