## Supplementary figures and images for "Interferon α and β induce differential transcriptional and functional metabolic phenotypes in human monocyte-derived macrophages and blunt glycolysis in response to antigenic stimuli"

### Supplemental figure S1

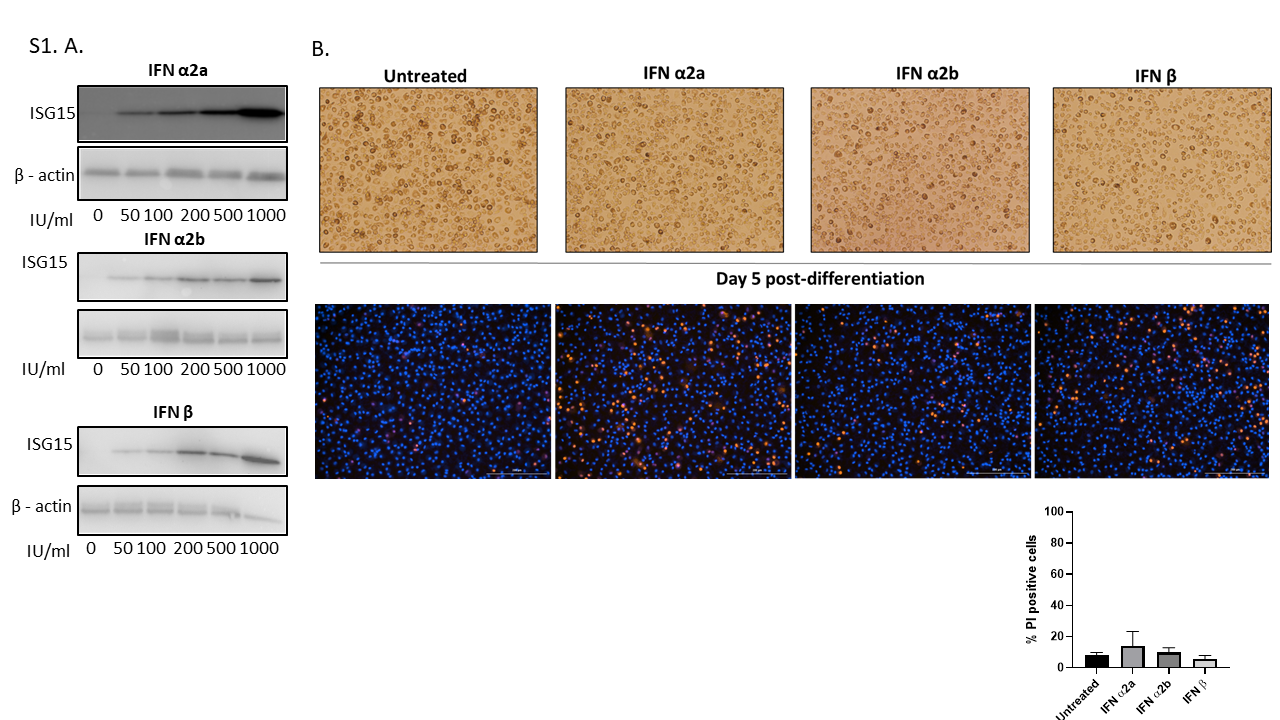
